# A Globally Representative Immunopeptidomics Approach to Identify Population-Wide Vaccine Candidates

**DOI:** 10.1101/2025.11.25.690604

**Authors:** Teesha Baker, Lucy Song, Charley Cai, Selwyn Gu, Leonard J. Foster

## Abstract

Vaccine development has historically relied on preclinical testing using one or two cell lines, typically of Caucasian descent, contributing to vaccines that may fail to provide adequate protection across genetically diverse populations. A major factor in these failures is the lack of consideration for human leukocyte antigen (HLA) diversity, with over 30,000 HLA alleles worldwide exhibiting distinct frequencies across ethnic groups and geographic regions. Here, we present a globally representative immunopeptidomics approach that addresses HLA diversity at the earliest stages of vaccine antigen discovery. We established a panel of 30 cell lines from the 1000 Genomes Project that represent 75% of the world’s most common HLA alleles. Using a combined computational and experimental approach, we identified 18 common binding motifs (7 MHC class I and 11 MHC class II) shared across 24 cell lines from the panel. We developed an accessible bioinformatic tool that predicts globally presentable antigenic regions from any pathogen proteome by identifying peptides matching these common binding motifs. Validation with *Salmonella enterica* serovar Typhimurium and SARS-CoV-2 Spike protein demonstrated that our method successfully identifies known immunogenic regions, with experimentally detected epitopes showing substantial overlap with predicted regions of interest. Notably, only three cell lines were required to validate highly immunogenic *S. enterica* targets, including OmpC—a porin with known 100% protective efficacy—detected across all tested lines. Our approach enables researchers to rapidly screen pathogen proteomes using freely available bioinformatic tools, with optional experimental validation requiring minimal resources. By identifying antigens with broad population coverage before clinical trials begin, this method has the potential to increase vaccine success rates while ensuring equitable protection across diverse populations. The cell line panel, common binding motifs, and bioinformatic workflow are publicly available, providing an accessible pathway for developing vaccines that truly serve global populations.

## Introduction

Vaccines traditionally include antigens either derived from the target pathogen or components produced synthetically to induce an immune response (1). This approach has led to the eradication or control of various pathogens, saving 3.5-5 million lives annually (2). Older vaccines use live or attenuated forms of the pathogen itself, while modern ones use subunit or single proteins from the infectious agent. Subunit vaccines are considered safer as they are unable to cause disease and are less taxing on the immune system (3,4), making them more suitable for immunocompromised individuals compared to full-pathogen vaccines. A key step in this process is identifying which protein(s) would be the most immunogenic thereby providing the greatest protection. For example, previous research into coronaviruses indicated that the Spike protein was a key antigenic target (5), so this protein or its mRNA transcript is used in SARS-CoV-2 vaccines. For most pathogens, however, the optimal antigens are less obvious.

The consequences of neglecting HLA diversity in vaccine development are increasingly evident. In COVID-19 vaccine clinical trials, 85% of participants were of Caucasian ethnicity, with only two studies reporting vaccine efficacy among ethnic minority groups (Salari et al. 2022). HLA polymorphism can complicate vaccine design and development, especially for vaccines containing selected epitopes directed at pathogens prevalent worldwide, as such vaccines may not be uniformly effective across populations due to HLA class I restricted antigen recognition and ethnic variation in HLA distribution (Dawson et al. 2001). Variation in human leukocyte antigen impacts the immune response, and genotypic variability in vaccine response may affect the safety and efficacy of potential vaccines (Pepperrell et al. 2021). The malaria vaccine RTS,S illustrates this challenge: while providing significant public health benefit, substantial heterogeneity was noted in efficacy between population subgroups, with vaccine efficacy against clinical malaria lowest at high transmission intensity (Bejon et al. 2013). Polymorphisms in T-cell regions can affect T-cell recognition and HLA binding, potentially compromising vaccine efficacy (Takala et al. 2013). These examples underscore the critical need to consider HLA diversity from the earliest stages of vaccine development rather than discovering limitations during clinical trials.

The original process for antigen identification was slow and empirical: limited candidates (∼10-25) would be chosen through a combination of biochemical and genetic methodologies and then tested *in vivo* for their immunogenic capacity (6), with few candidates passing clinical trials. Reverse vaccinology is a technique that uses computation tools to predict antigenic regions from the pathogen’s genome. Full or partial versions of candidate proteins numbering in the hundreds or thousands are then made synthetically or recombinantly to test for their immunogenicity *in vivo* (7). This method was first proposed in 2000 to identify vaccine candidates from the *Neisseria meningitidis* serogroup B genome (8). Since then, reverse vaccinology has successfully created vaccines for *Klebsiella pneumoniae* (Dar *et al*., 2019), *Staphylococcus aureus* (Kim *et al*., 2011), *Leptospira interrogans* (Gamberini *et al*., 2005), and *Acinetobacter baumannii* (Singh, Capalash and Sharma, 2017).

Another antigen discovery approach, immunopeptidomics, detects pathogen-derived peptides presented by major histocompatibility complex (MHC) molecules using liquid chromatography coupled with tandem mass spectrometry (LC-MS/MS). The conventional method for obtaining immunopeptides by immunoprecipitation (IP) (Bassani-Sternberg *et al*., 2016) relies on antibodies and does not scale well across MHC alleles. Mild acid elution (MAE) is an alternative method that simply and quickly elutes peptides presented on the cell surface for presentation to T-lymphocytes (Storkus *et al*., 1993, Chan et al, 2025). The omission of antibodies allows the inclusion of weaker binding ligands (Sturm *et al*., 2020). Furthermore, IP exhibits bias against MHC-bound peptides that are highly hydrophobic or hydrophilic (Sabdia *et al*., 2022), leading to the loss of some potentially highly immunogenic ligands. For these reasons, we selected MAE as our method for experimental validation of computationally predicted binding motifs, using biological replicates and stringent filtering criteria to ensure data quality.

Despite many advances in vaccine development, one major hurdle is that many antigens that appear promising in pre-clinical studies may fail to show the same efficacy in humans, particularly across genetically diverse populations (Rando *et al*., 2023). Generally, small-scale studies limited to a single geographical area show promise but drop in efficacy when transitioning to broader, more heterogeneous populations (Wong, Siah and Lo, 2019). One of the significant factors contributing to the failure of clinical trials is the lack of consideration for the genetic diversity of human leukocyte antigens (HLAs) in the target population. HLA genes are highly polymorphic, with over 30,000 alleles worldwide (Robinson *et al*., 2020). Different ethnic groups from various geographic areas exhibit distinct HLA haplotype frequencies (Cao *et al*., 2004; Guech-Ongey *et al*., 2010; Yao *et al*., 2012). A good vaccine must contain antigens that can be presented on many (if not most) HLA alleles to generate immunological memory and protection across the entire population. However, this is difficult to achieve as an antigen that is strongly-presented by one MHC allele may not be equally presentable by other alleles, leading to variable immune responses. This highlights the importance considering MHC diversity at the onset of vaccine design across different ethnicities and geographical regions. Considering the global frequency and distribution of HLA polymorphisms during both the discovery and testing phase will increase the success rate of vaccines at a global scale. More broadly immunogenic vaccines, fully immunizing a greater percentage of those who receive the vaccine, can be created by using *rational vaccine design*; a set of highly immunogenic vaccine candidates can be identified through experiments utilizing the naturally occurring mechanisms for protein processing and subsequent antigen presentation, while incorporating allelic diversity with global representation in a cell line panel with multiple HLA haplotypes.

To address this issue, we introduce an innovative approach to identify potential antigens for vaccine development by comparing immunopeptide profiles in cell lines representing the wide genetic diversity found in the HLA genes worldwide. The cell line panel outlined herein offers the potential for in-house testing and antigen selection for many diverse pathogens in globally representative pre-clinical experiments. We aimed to cover as large a fraction of the world population as possible, while ensuring utility for researchers, by using a publicly available collection of immortalized antigen-presenting cell lines that offer comprehensive global genetic representation (Auton, 2015). This approach offers the advantage of speed while simultaneously accounting for high genetic diversity, providing a novel pathway for vaccine development that better addresses the challenges posed by global genetic variations.

## Materials & Methods

### Selection of Cell Lines

Cell lines used for this project were selected from the 1000 Genomes Project (https://www.internationalgenome.org/), an international genome sample resource that has aimed to create a deep catalog of human genetic variation (Auton, 2015). There are over 3000 unique patient EBV-transformed B-lymphoblastoid cell lines (BLCLs) available in the database. Haplotype and geographical location data for the 3000+ cell lines available for purchase by the 1000 Genomes Project were downloaded from the FTP server, http://ftp.1000genomes.ebi.ac.uk/vol1/ftp/. The 1000 Genomes Project provides documentation for up to two heterozygous alleles for each of the following MHC genes: HLAs A, B, C, DQB1, and DRB1 (missing the α assignment of DP and DQ, and C entirely). An in-house algorithm was used to rank and select cell lines from the 3000+ cell lines available in order to maximize for global HLA-DR diversity in the lowest number of cell lines. Ranking was performed by calculating the frequency of each allele that occurred within the database from all cell lines. Each cell line could then be ranked against all others in the database for the coverage that that cell line represents within the 3000 cell lines, with the cell lines with the most commonly occurring HLAs, or most representative cell lines, ranked at the top. The selected BLCLs were purchased from Coriell Institute for Medical Research. A list of BLCLs used in the study and their HLA haplotypes are included in Supplementary Materials.

### Haplotyping with Exome Data

Data from the 1000 Genomes Project can be viewed in multiple formats at https://www.internationalgenome.org/data. All data files are available for download on their FTP server (link above). For each of the 30 cell lines, the 30× whole-genome BAM file was downloaded. Using xHLA2, the full allele set for class I alleles (HLA-A, HLA-B, and HLA-C) and class II alleles (HLA-DPA, HLA-DPB, HLA-DQA, HLA-DQB, and HLA-DRB1/3/4/5) were assigned. These alleles can be found in the Appendix - Table S1, Table S2, and Table S3.

### Culturing Mammalian cells

BLCLs were purchased and shipped in non-vent cap T-25 flasks. Upon receiving, cultures were allowed to equilibrate back to incubator conditions for 24-h before counting. Cells were cultured in Roswell Park Memorial Institute (RPMI) 1640 media with 1x L-glutamine (Gibco), 10% FBS (Gibco), and 1% penicillin-streptomycin (Gibco). Cells were maintained in an incubator (PHCbi) at 37°C with 5% CO_2_. To sustain optimal cell health, media were replenished every other day when the cell concentration reached 1x106 cells/mL, with cell viability >75%. Fresh media was then added to the culture, without removing conditioned media, and adjusted accordingly to achieve the desired cell concentration of 5x105 cells/mL. In cases where cell concentrations dropped below the desired range and/or viability fell below 75%, cells underwent a centrifugation step using the Fisher Scientific AccuSpin 1R Refrigerated Centrifuge at 90 x g for 5 min to remove cellular debris. Following centrifugation, the supernatant was carefully aspirated, and cells were resuspended in 100% fresh media to attain the desired concentration. Accurate cell counts were performed using Trypan Blue staining (0.4%, Invitrogen) in conjunction with an automated cell counter (Thermo Fisher Countess II). The Trypan Blue solution was mixed with the cell suspension at a 1:1 ratio and loaded onto a hemocytometer for fast and precise cell counting.

### Culturing Bacteria Cells

*Salmonella enterica* (SL1344) were streaked with an inoculation loop into lysogeny broth (LB plate: 1L: 10 g tryptone, 5 g yeast extract, 10 g NaCl, 1.5% agar, LB media: 10 g tryptone, 5 g yeast extract, 10 g NaCl) plates with 0.01% streptomycin. Select colonies were cultured in spinner flasks with 1L LB and 0.01% streptomycin overnight in a shaking incubator (37°C, 140 rpm, New Brunswick Innova 42/42R Stackable Incubator Shaker).

### Pathogen exposure

Bacteria cells were lysed in 1x PBS using a 60 minute autoclave run.BLCLs were individually exposed with SL1334 at a ratio of 10 mg bacteria lysate to 5x10^7^ mammalian cells (N=63 per cell line). Cells were harvested for antigen collection by mild acid elution, using the wash and elution protocol details in the next methods section, after being exposed for 24-h at normal growing conditions (37°C, 5% CO2).

### Mild Acid Elution

Prior to the mild acid elution procedure, cell counts and viability were assessed to ensure suitable conditions for antigen extraction. A total of 5x107 cells were required for each repeated biological replicate (N=6 for non-exposed and N=3 for bacteria lysate exposed), with a minimum viability threshold of 85%.Cells were pelleted at 500 x g for 3 min (Sorvall T1 centrifuge, Thermo Scientific) in 50mL conical tubes. The supernatant were discarded and the pellets were washed once with 10mL phosphate buffered saline (PBS, 1x, then twice with 10mL cold PBS. The washed pellet was then resuspended in 10mL cold saline (PBS without either of the cations’ phosphate salts, 1x) and transferred to fresh 50 mL tubes to remove residual phosphates. Cells were pelleted again under the same conditions, discarding the supernatant. Cell surface presented antigens were eluted by resuspending cells with 2% acetic acid in the same 1X cold saline as before. After a 30 second exposure to the acid solution, cells were pelleted by centrifugation and the supernatants were collected. Samples were either frozenat −80°C overnight or flash frozen in liquid nitrogen then placed in a lyophilizer (Labconco) for 48 h. The dried powder was stored in −80°C until use.

### Offline Liquid Chromatography Fractionation & Subsequent Concatenation

Lyophilized samples were resuspended in Buffer A and desalted using STAGE tips as indicated in Rappsilber, Ishihama and Mann, 2003. Desalted samples were resuspended in Buffer A then fractionated using an Agilent ZORBAX Extend column (80 Å C18, 1.0 x 50 mm, 3.5 μm) on a high-performance liquid chromatography system (Agilent, 1200 Series) with a 36 minute gradient of Buffer A (2% acetonitrile in 5 mM NH4HCO2, pH 10) and Buffer B (90% acetonitrile in 5 mM NH4HCO2, pH 10). The gradient parameters were as follows: 0 to 4% Buffer B for 30 s, increased to 40% over 20 min, then held at 90% for 5 min, followed by re-equilibrated for the remaining 10.5 min at 0% B. Fractionation produced 12fractions which were subsequently concatenated into four by combining fractions 1, 5, and 9, fractions 2, 6, and 10, fractions 3, 7, and 11, and fractions 4, 8, and 12 together. Concatenated samples were dried by centrifuge concentrator (1400 rpm, Vacufuge plus, Eppendorf) and desalted again by STAGE tips as described above and dried by vacuum centrifugation.

### Online Liquid Chromatography & Mass Spectrometry

Desalted and dried samples derived from MAE were reconstituted in 20 μL of Loading Buffer (2% acetonitrile in 0.1% formic acid). A NanoDrop One (ThermoFisher Scientific) was used to quantify (by Scopes method by measuring absorbance at 205 nm) peptide concentration. Samples were diluted and 50 ng were injected into the Easy nano LC 1000 HPLC (ThermoFisher Scientific) coupled to a timsTOF Pro (Bruker Daltonics) via Captive Spray nanospray ionization source (Bruker Daltonics). Reverse phase chromatography was carried out using an Aurora Series Gen2 (25 cm x 75 μm with 1.6 μm C18 beads, a 120 Å pore size, with Gen2 nanoZero and CSI fitting by Ion Opticks (Parkville, Victoria, Australia). The column was heated to 50°C using a tape heater (SRMU020124, Omega), and controlled using an in-house microprocessor. Samples were separated at a flow rate of 0.35 μL/min with a 45 min gradient ; 5 to 30% Buffer B (90 % acetonitrile in 0.1% formic acid) over 45 min, then increased to 100% Buffer B over 2 min, and held at 100% Buffer B for 13 min. Peptides derived from full cell lysates were separated on a 90 min gradient: 5% to 15% Buffer B over 45 min, then increasing to 30% Buffer B over the next 45 minutes, then to 100% B over 2 min, and finally held at 100% B for 13 min. The timsTOF was set to acquire in a data-dependent parallel accumulation-serial fragmentation (PASEF) mode,fragmenting the 10 most abundant ions, including +1 ions by drawing the ion mobility zone of interest to include +1 ions (one at the time at 18 Hz rate) after each full-range scan from 100 to 1700 Th. The nano ESI source was operated at 1900 V capillary voltage, 3 L/min drying gas and 180°C drying temperature. Funnel 1 was set at 300 V, funnel 2 at 200 V, multipole RF at 200 V, deflection delta at 70 V, quadrupole ion energy at 5 eV, low mass at 200 Th, collision cell energy at 10 eV, collision RF at 1500 V, transfer time at 60 μs, and pre-pulse storage at 12 μs. PASEF was on with 10 PASEF scans for charges 0 to 4, Target intensity 20000 and Intensity threshold 2500.

### Peptide Spectrum Matching

Fragment spectra matching for peptide identification was performed with FragPipe (Kong *et al*., 2017). All MAE-derived samples were searched against human and bovine protein entries from Uniprot and SwissProt (annotated and reviewed) entries, as well as all Uniprot, SwissProt, and TrEMBL *S.enterica* (serovar Typhimurium) entries with decoys generated by Fragpipe. The search was conducted with no enzyme specificity with peptide lengths limited to7-25 amino acids. Other parameters included precursor mass accuracy of 40 ppm and fragment mass accuracy of 20 ppm. Variable methionine oxidation was included and a 1% false-discovery rate (FDR) cut-off was used at the protein and peptide level.

### Data Analysis

All code used to perform data analysis can be found on GitHub at https://github.com/teeshabaker/MHC_MAE. Full lists of identified peptides were loaded into R statistical software (version 4.3.1, R Core Team) using the RStudio IDE (version 2021.9.0.351, RStudio Team) for data wrangling and manipulation. The first step was to perform basic filtering, which removed contaminants, duplicates, and reverse peptides. All peptide IDs were kept when combining fractions from the same repeated measure (but may be fractionated into multiple MS injections), then when combining repeats for a single list of peptides from a given cell line, the peptide entry must be present in at least two out of three replicates to be considered a positive peptide identification. After filtering, a length cut-off was applied to all peptides to create two groups: one with peptides between 9 to 11 amino acids long, and a second group with peptides between 12 to 20 amino acids. Next, peptides in each group were compressed into epitopes using an in-house script to group overlapping sequences together, outputting a single sequence of the union of the group.

The full haplotype of each cell line is required for analysis and interpretation of the results of immunopeptidomic experiments. The exome data for the panel of the 30 chosen cell lines from the 1000 Genomes Project were downloaded for HLA allele assignment with xHLA (Auton, 2015; Xie et al. 2017). Most of the new HLA assignments were the same as the original allele assignments for the 30 cell lines, however, in any cases where there was a discrepancy, the new assignment from xHLA was used (Xie et al. 2017).

Each peptide was assessed for MHC binding by using netMHCpan for peptides length 9 to 11 amino acids or netMHCIIpan for peptides length 12 to 20 amino acids to evaluate each peptide against a set of naturally eluted ligands to predict the binding affinity and core region of that peptide (Reynisson *et al*., 2020). Cores of 9 amino acids were searched against the specific MHC I and MHC II alleles from that cell line’s haplotype. For netMHCpan, peptides with a score of 0.5% or less were considered a strong binder and 0.5 to 2% were considered a weak binder. For netMHCIIpan, peptides with a score of 1% or less were considered a strong binder and between 1 and 5% were considered a weak binder. For an epitope to be considered a binder, only one binding core needed to be present in the union sequence. For epitopes that were predicted to bind to multiple HLAs, the strongest binding prediction value (closest to 0%) was used to assign its HLA. All epitopes, regardless of their binding prediction value, were submitted to GibbsCluster for alignment and grouping of sequences (Andreatta, Alvarez and Nielsen, 2017). Seq2Logo was used to create sequence logos from the alignments derived from each cluster (Thomsen and Nielsen, 2021).

## Results

### Cell line selection for genetic and geographical diversity

In order to develop a methodology for identifying antigenic peptides that provide global population coverage, we needed to establish a representative cell line panel that contains HLA alleles expressed across all major populations across the globe. Certain alleles are common across all continents and geographical regions, while others are more unique to small regions. We took advantage of the genetic data and cell lines available from the 1000 Genomes Project whichwas curated with the goal to capture this genetic diversity among and within populations across the globe.(Auton, 2015). It was unknown how many cell lines would be required to account for the global genetic diversity within HLA alleles. To determine an appropriate number of cell lines required, the HLA allelic information from the 3000 cell lines provided was used as a baseline to calculate each allele’s frequency within the database.

We based our allele frequency calculations on HLA-DRB1, the most diverse HLA gene, but this pattern was consistent across all alleles, confirming there is a comprehensive representation of HLA allelic diversity within the chosen cell line panel. We ranked each of the 3000 cell lines for their ability to represent as many of the global HLA-DRB1 alleles as possible, with a higher rank given to those that expressed a higher frequency haplotype. The ranking results can be seen in Figure 1a and b). The plateau points of these data indicate the number of cell lines at which including more would no longer increase the overall total genetic coverage within the selected cell lines. This saturation of coverage was achieved at approximately 30 cell lines. Based on these data this is the minimal number of cell lines that should be included in in vivo pre-clinical trials involving the immune system and MHC genes for minimum global representation. The top 30 cell lines from this ranking process created our representative pre-clinical trial panel and can be found in Supplementary Table S1-3. Due to financial constraints, 14 of the 30 cell lines in the representative panel were chosen at random for culturing and immunopeptide analysis. While the COVID-19 pandemic highlighted the importance of diverse representation in clinical trial enrollment, with vaccine trials achieving greater ethnic diversity than historical standards, the fundamental question of whether selected vaccine antigens can be presented across diverse HLA alleles has remained largely unaddressed in the preclinical phase. Our approach tackles this issue at its source by ensuring antigenic regions are selected for their potential to be broadly immunogenic before any clinical trials begin.

**Figure 1.**
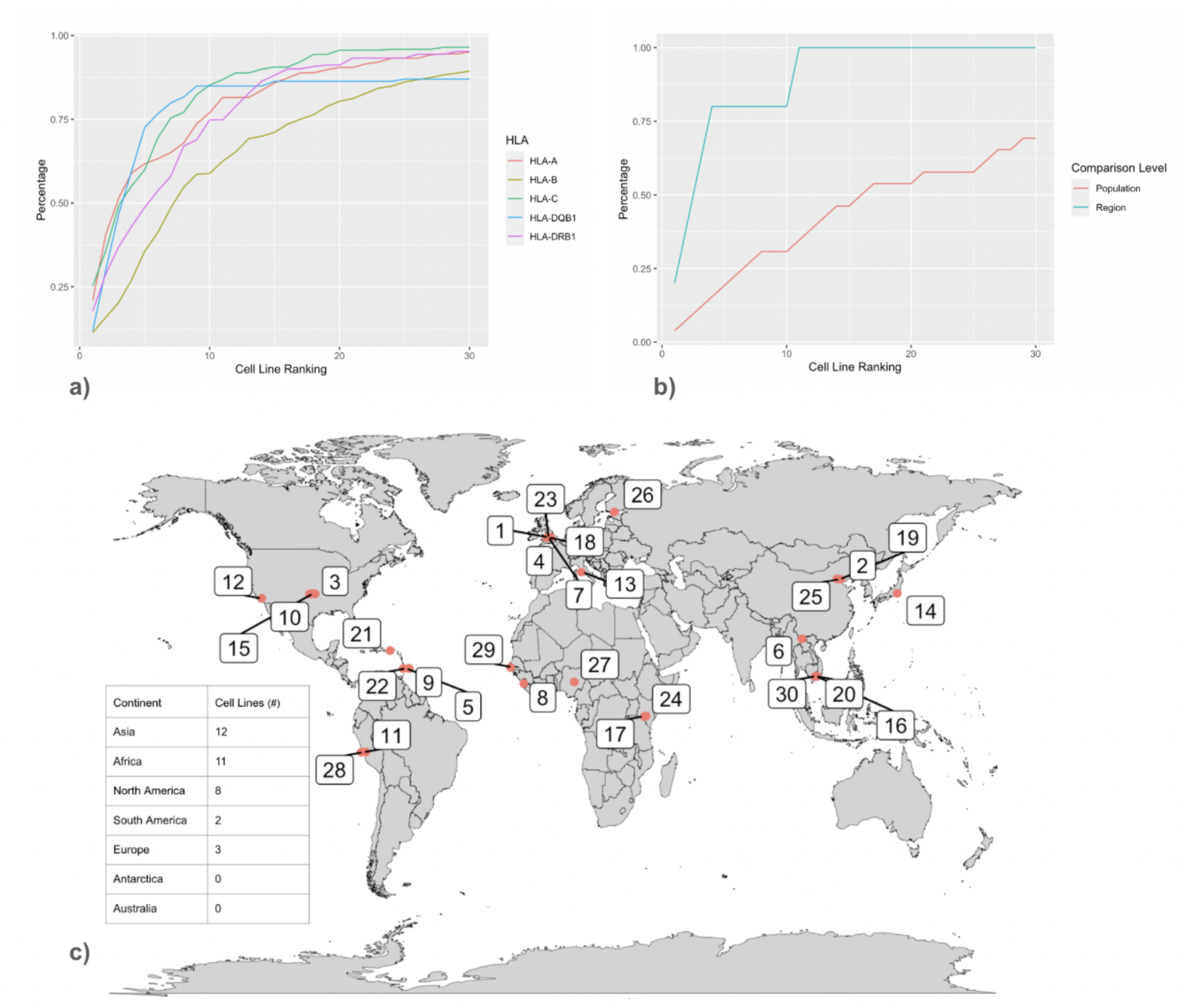
1000 Genomes Project Alleles (Auton, 2015). The cumulative coverage of HLA alleles in the 1000 Genomes Project (Auton, 2015) a) The cumulative coverage of all HLA alleles of the global population sampled by the 1000 Genomes Project (Auton, 2015) from their list of cell lines ranked based on the frequency of the provided HLA-DRB1 allele within the database. b) The cumulative coverage of the number of regions and populations present from their list of cell lines ranked based on the frequency of HLA-DRB1 allele within the database. c) The geographical origins of the 30 selected cell lines. The numbering corresponds to the cell lines listed from top to bottom in Table S1-3.

It was critical to select a number of cell lines that ensured adequate spread across the “regions” and “populations” (as defined by the 1000 Genomes Project) while maximizing geographical representation. The database has a total of 31 populations, however, only 26 of the 31 contain an immortalized cell line + NGS data required for all experiments. While this selection process does not explicitly take into account geographical regions or populations selected, these cell lines appear evenly distributed across the globe (Figure 1c) and represent 70% (19 out of 26, Figure 1b) of the populations available in the 1000 Genomes Project (Auton, 2015). Increasing the cut-off in the ranked allele frequency list up to 50 cell lines without targeting specific populations or alleles achieved only 85% coverage of the total populations (not shown). For any future experiments that want to increase HLA coverage, it would be advised to manually select additional cell lines from the ranked list to target specific populations or HLA alleles, rather than increasing the cut-off cell lines selected from the list. This panel represents a paradigm shift from conventional vaccine development, which typically uses only one or two cell lines for preclinical characterization. By incorporating data from cell lines representing 70% of global populations and 75% of the world’s most common HLA alleles into early-stage antigen discovery, our approach ensures that vaccine candidates are screened for broad population coverage before significant resources are invested in candidates that may have limited global efficacy.

The haplotype information provided online by the 1000 Genome Project was incomplete; HLA-C and the α assignment of HLA-DP and HLA-DQ is not available online. This missing data did not cause any apparent issues with selecting a diverse panel of cells; the plateau points of all alleles tested were at ∼30 cells (Figure 1a) and cells were geographically distributed (Figure 1c). However, the full haplotype of each cell line is required for analysis and interpretation of the results of immunopeptidomic experiments. (e.g., netMHCIIpan requires both the α and β chain assignments for prediction). The whole genome sequencing and full exome data for all 3000+ cell lines are freely available for download from https://www.internationalgenome.org/data. NextGeneration Sequencing (NGS) provides access to much larger multiplexed and untargeted datasets, however, these need to be aligned in order to interpret the data. xHLA is a command line script algorithm written for HLA typing based on translated short reads. It uses an exhaustive multiple sequence alignment-based expansion, and iterative solution set refinement at the amino acid level to determine the full allele set at four-digit typing accuracy, including DRB3/4/5 if available (Xie, C. et al. Fast and accurate HLA typing from short-read next-generation sequence data with xHLA. Proc. Natl. Acad. Sci. 114, 8059–8064 (2017)). The exome data for the panel of the 30 cell lines downloaded for HLA allele assignment with xHLA. The full allele set (16 alleles total, heterozygous assignments for class I alleles (HLA-A, HLA-B, and HLA-C) and class II alleles (HLA-DP, HLA-DQ, and HLA-DR), with variations in DP and DQ occurring within both the α and β chains) and optional DRB3/4/5 were obtained, which can be found in the Supplementary (Table S1-3). Most of the new HLA assignments were the same as the original allele assignments for the 30 cell lines, however, in any cases where there was a discrepancy, the new assignment from xHLA was used. HLA-C, HLA-DPA, and HLA-DQA were all new from xHLA. These data represent a now complete allele set, which can be used to search against software like netMHCpan that was missing the HLA-DP and DQ alleles entirely when searching for predicted binders. In addition, DRB3/4/5 were missing from the original haplotype information, yet contribute significantly to the total DR immunopeptidome (Kaabinejadian, S. et al. Accurate MHC Motif Deconvolution of Immunopeptidomics Data Reveals a Significant Contribution of DRB3, 4 and 5 to the Total DR Immunopeptidome. Front. Immunol. 13, 835454 (2022)). With full haplotype information, the selected cell lines were compared to the global allele frequency for a comparison of true global coverage. Based on the top 42 alleles in the Allele Frequency Net Database (Gonzalez-Galarza et al. 2020), 75% of these most common alleles are represented at least once in our panel. This panel provides a valuable resource for future investigations in pre-clinical vaccine development and genetic diversity studies.

### Identification of common binding motifs

Our goal was to identify common binding motifs (CBMs) that were shared among our panel of 30 cell lines. This was achievable by combining the data from two methods: bioinformatics (dry bench) and MAE (wet bench). While it is possible to find CBMs using either one of the two methods alone, the data was significantly improved by using both. Here we will first discuss the bioinformatic methodology.

Of the 30 cell lines, we selected the top 15 for analysis, which already covered over 85% of the HLA-DR alleles available in the 1000 Genomes Project (Auton, 2015, Figure S3). Their binding motifs were found on MHC Motif Atlas (Tadros *et al*., 2023). For each motif that was common to at least five of our cell lines, their relevant ligand sequences were downloaded, clustered, and aligned using GibbsCluster (Andreatta, Alvarez and Nielsen, 2017) to produce logo plots. In total, 7 MHC I and 9 MHC II CBMs were found, and 3 were discarded due to insufficient data (Figure S1).

To supplement these results, wet bench experiments using MAE were done for 14 of the 30 cell lines (Figure S3). Identified sequences were separated by length (9-12, and 13+ for MHC-I and MHC-II respectively), collapsed into epitopes by grouping overlapping sequences, and then submitted to netMHCpan to determine their binding strength. For netMHCpan, peptides with a score of 0.5% or less were considered to be strong binders and 0.5 to 2% were considered weak binders. For netMHCIIpan, peptides with a score of 1% or less were considered a strong binder and between 1 and 5% were considered a weak binder. For an epitope to be considered a binder only one binding core needed to be present in the union sequence. For epitopes which were predicted to bind to multiple HLAs, the strongest binding prediction value (closer to zero %) was used to assign its HLA. All epitopes predicted to bind HLA were submitted to GibbsCluster for the alignment and grouping of peptides. Motifs were considered CBMs if they were identified more than once. The sequences were again clustered and aligned by GibbsCluster, which resulted in a total of 7 MHC I and 3 MHC II CBMs (Figure S2).

To combine data generated from bioinformatics and MAE, CBMs that were found by both methods were “regenerated” by pooling their ligand sequences and resubmitting them to GibbsCluster. No further treatments were necessary for the CBMs that were discovered in only one of the two dry- and wet-bench techniques. In total, 7 MHC I and 11 MHC II CBMs were found (Figure 3). This covers 24 of the 30 selected cell lines that represent 60% of the global populations (Figure 1). As a result, antigens that contain these motifs are more likely to make suitable antigens for vaccines.

**Figure 3.**
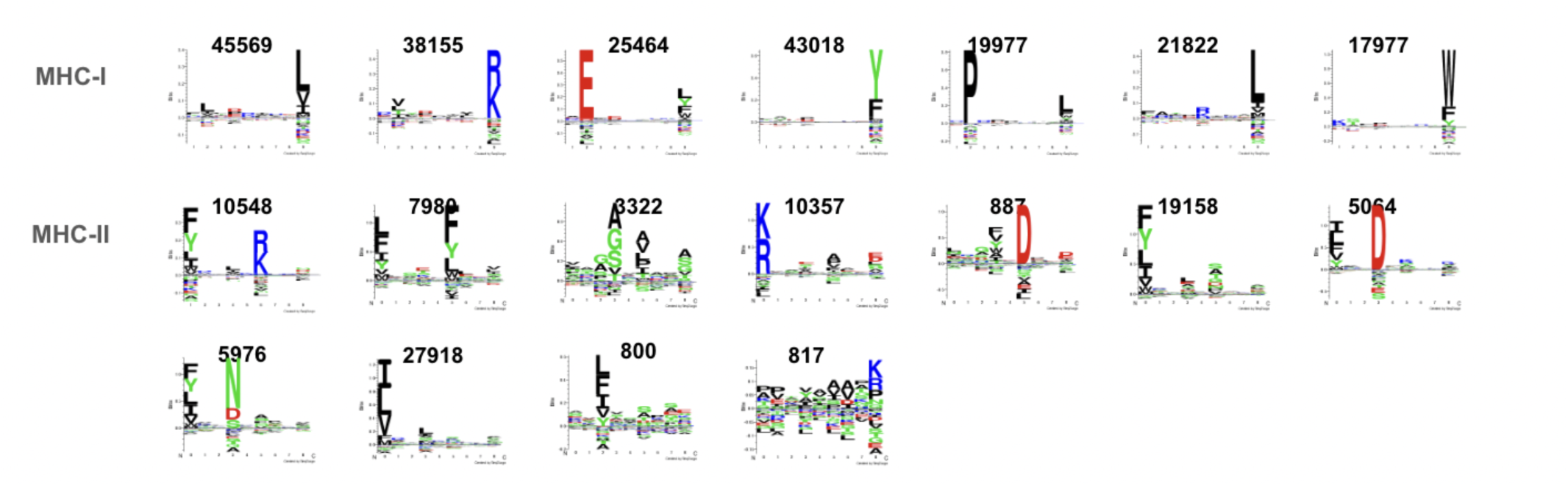
Logo plots of common binding motifs generated from peptides derived from the MHC Motif Atlas (Tadros *et al*., 2023) combined with our MAE-derived sequences. Common binding motifs were identified from a total of twenty four cell lines. The numbers above the motif indicate the total number of unique peptides used for generating the sequence logo.

**Figure 4.**
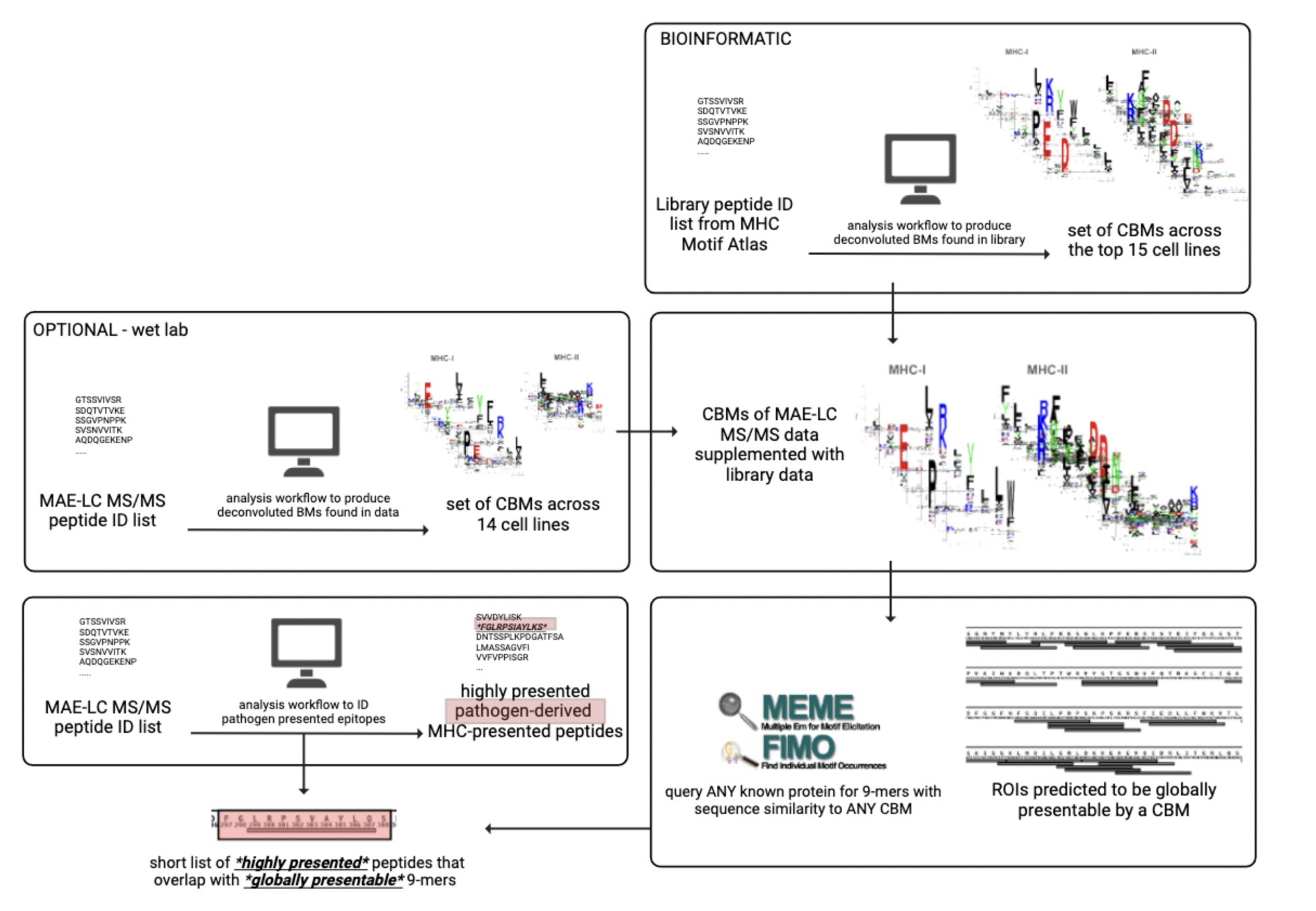
The construction of our workflow. Data from bioinformatic analysis and optional wet-bench procedure can be used and combined to create CBMs. These motifs can help find regions of interest (ROI) from a pathogen’s genome or proteome sequences that are likely to be presented on MHC proteins of a large portion of the human population.

### Bioinformatic tool to identify globally presentable peptides

Using our data, we aimed to develop a tool that identifies protein regions from pathogens that are likely to be presented on the MHCs of a significant portion of individuals around the globe. All 18 CBMs we identified were represented by matrices incorporating the anchor positions within the binding cleft, notably positions 2 and 9 which are essential for binding (Tirado-Herranz et al. 2023; Mukherjee et al. 2021; Cresswell et al. 2005; Roche and Furuta 2015). To do this, we utilized the MEME Suite (Bailey, Johnson, and Grant, 2015), which is a set of tools designed to create, query, and manipulate sequence matrices from both DNA and protein data. One of the softwares in the suite, Find Individual Motif Occurrences (FIMO) (Grant, Bailey and Noble, 2011) can scan a target protein for any occurrences of a given motif. This software was used to produce a shortlist of peptides that contain CBMs from various pathogen proteins that are meant to be tested in various immunogenicity assays such as HLA-matched T-lymphocytes in an enzyme-linked immunospot (ELISPOT) sandwich enzyme-linked immunosorbent (ELISA) assay. Our tool can be used/downloaded at https://github.com/teeshabaker/MHC_MAE.

### Testing our workflow on a common pathogen

We tested the ability of our workflow to identify ROIs on *S. enterica* serovar Typhimurium against experimentally detected MHC-bound peptides from exposed cells. Three cell lines were exposed to *S. enterica* lysate for 24 h, then host cell surface peptides were collected by MAE. A total of 19 different *S. enterica* epitopes were detected (Table S4). There was only a single *S. enterica* epitope that was detected in all three cell lines: OmpC. From one of the cell lines (GM20346), we identified two more epitopes from Omp D and the 30S ribosomal protein S7. All three were part of nested peptide sets, which is indicative of highly immunogenic antigens (Geysen et al., 1984)(Figure 6). Exposure to Omps have been shown to protect the host against lethal doses of *S. enterica* serovar Typhimurium with 100% efficacy in one study (Hamad *et al*., 2008). In a different experiment with *S. enterica* serovar *Typhi* (Pérez-Toledo et al., 2017), OmpC and OmpD are both porins that are highly immunogenic. We compared the three experimentally identified epitopes of *S. enterica* serovar Typhimurium against the ROIs found by our workflow. All of them showed a sizable overlap to the bioinformatically derived ROIs, implying that they are likely to be suitable vaccine candidates that will be immunogenic for a substantial portion of the global population.

These findings are particularly significant given that only three cell lines were required to identify these highly immunogenic targets, demonstrating the efficiency of our approach. The detection of OmpC across all three genetically diverse cell lines, combined with its known 100% protective efficacy against lethal S. enterica serovar Typhimurium challenge (Hamad et al., 2008), validates that our method successfully identifies antigens with both broad HLA presentation and protective potential. This experimental validation required minimal resources compared to traditional approaches, yet provided strong evidence for global immunogenicity.

### Testing our workflow on a known antigenic protein

We also tested our bioinformatic tool on the SARS-CoV-2 Spike protein to find out if the ROIs that we identified match known antigenic regions from existing literature. These highly presented S protein epitopes were from a recent clinical trial, BNT162b4 (Arieta *et al*., 2023), which successfully increased the efficiency of the Pfizer- BioNTech COVID-19 Vaccine (BNT162b2) (Lamb, 2021) with an additional mRNA component with sequences identified from an HLA-I immunopeptidomics experiment (Weingarten-Gabbay *et al*., 2021).

Immunoprecipitation (IP)-LC-MS/MS identified pathogen-derived MHC-presented immunopeptides from SARS-CoV-2 infected cells. Data from this experiment and their HLA-II experiment (Weingarten-Gabbay *et al*., 2024) were utilized. Most of the presented peptides identified occur in a region of the protein with multiple presentable 9-mers on one or multiple of the common binding motifs (Figure 7). Not every presentable ROI will be seen in this dataset, as not all common binding motifs would occur in the two cell lines used for infection (A549, a lung epithelial cell line; and HEK293, a kidney cell line)(Weingarten-Gabbay *et al*., 2021; Weingarten-Gabbay *et al*., 2024). The overlap between our globally presentable predictions and known antigenic regions identified through multiple independent studies demonstrates that our method captures functionally relevant epitopes. Importantly, this analysis was performed entirely computationally using our published common binding motifs—requiring no wet-bench work whatsoever—yet successfully predicted regions that are independently validated as immunogenic. This illustrates the practical accessibility of our approach: researchers can rapidly screen pathogen proteomes for globally presentable regions using only bioinformatic tools, reserving experimental validation for the most promising candidates. Beyond the immunopeptidomics data, our predictions also aligned with structurally defined antigenic regions.

Additionally, other regions of the S protein that have been identified as antigenic in literature were overlaid (striped coloured bars). These regions also have a high percentage of overlap with *globally presentable* epitopes. A recently identified putative superantigen (Cheng *et al*., 2020), though only 22 AAs, has two overlapping 9-mers predicted to bind one of the common binding motifs. The receptor-binding domain, which is a highly targeted region of the S protein due to its interaction with the host binding receptor ACE2 (angiotensin converting enzyme-2), has the lowest density of presentable regions. The fusion peptide is centered on three overlapping 9-mers, as a small 18 AA peptide. The heptad repeats (HR1 and HR2) are also of interest, with the tail of HR1 a densely *globally presentable* region and HR1 overlapping 5 presentable 9-mers. A conformational change in a HR1 facilitates entry fusion peptide forward into the target membrane (Jackson, Farzan, Chen and Chloe, 2022; Cai *et al*., 2020). All of these regions could be tested individually for the efficacy as a subunit or mRNA-LNP vaccine, with the potential to include less non-presentable peptides that are not known to stimulate the immune system.

## Discussion

### Addressing a Critical Gap in Vaccine Development

A major hurdle in vaccine development is understanding which antigens will trigger an immune response in diverse populations, due in major part to the highly polymorphic HLA gene and differences in the sequences that the MHC proteins can present to T-lymphocytes. Current vaccine development practices exacerbate this problem by relying predominantly on cell lines of Caucasian descent during preclinical testing. This approach creates vaccines that may work well for certain populations but fail to provide adequate protection globally. The consequences of this narrow approach become apparent only after significant resources have been invested: clinical trials often fail when tested on genetically diverse populations, yet HLA diversity is rarely considered until that failure point. Our approach addresses this critical gap at the earliest stage of vaccine development—before significant resources are committed to candidates that may have limited global applicability.

The need for this paradigm shift is urgent. Vaccine clinical trials have historically suffered from poor ethnic diversity in participant enrollment, with minority populations underrepresented despite bearing disproportionate disease burdens. While COVID-19 vaccine development made strides in diversifying clinical trial enrollment, the underlying immunological question of whether vaccine antigens can be presented across diverse HLA alleles has remained largely unexplored in the preclinical phase. By the time diverse populations are enrolled in Phase II or III trials, the antigenic targets have already been selected—often without consideration for whether those targets will be immunogenic across different HLA backgrounds. Our method enables researchers to identify antigens with broad immunogenic potential before entering expensive and time-consuming clinical trials, thereby increasing the probability of global vaccine success.

### Methodological Innovation: Combining Computational and Experimental Approaches

Antigen discovery can identify numerous potential candidates, but defining which sequences will be presented by which HLA allele(s) is essential for triggering an immune response. It is not feasible to test every potential candidate on large and diverse populations during early development. While studies have investigated MHC I binding motifs to better understand binding preferences (Bassani-Sternberg et al., 2017), fewer studies have focused on MHC II-eluted peptides and their immunogenicity (Paul *et al*., 2019). This is validated by the lack of ligands available for some HLA class II alleles when generating our initial binding motif analyses (Figure S1). Notably, researchers have discovered that many HLA molecules can be grouped into fewer supertypes based on overlapping peptide-binding repertoires (Sette and Sidney 1998), and this HLA supertype-based strategy has gained validation in various disease settings (Sette and Rappuoli, 2020). Although numerous studies have been done on HLA class I classifications, further research is needed to fully characterize HLA class II supertypes (Sette and Rappuoli, 2020; Doytchinova and Flower 2005).

Similarly, while neural networks have become powerful tools for predicting antigen binding to MHC molecules and predictions for MHC-I are well established, reliable and consistent MHC-II binding predictions remain challenging (Sette and Rappuoli, 2020). The difference in predictive accuracy is largely attributed to the distinct ways these MHC classes bind peptide ligands. As such, some studies have explored using a consensus of many prediction methods (Sette and Rappuoli, 2020). Given these challenges, our project adopts a combined approach that integrates the identification of binding motifs through both a public database library method and experimental analysis of cell surface MHC-presented peptides using MAE. As demonstrated in our results (Figure S1 and Figures 2-3), although different panels of alleles were used, the HLA class I motifs remain consistent across both computational and experimental methods, while HLA class II motifs exhibit more variation— highlighting the value of our dual approach.

The MAE method offers several advantages over traditional immunoprecipitation (IP) approaches. By omitting antibodies, MAE includes weaker binding ligands and avoids bias against MHC-bound peptides that are highly hydrophobic or hydrophilic, leading to capture of potentially highly immunogenic ligands that IP would miss. While MAE can capture non-presented peptides from lysed cells or extracellular matrix our stringent experimental design addresses these concerns. The use of biological replicates (N=6 for non-exposed and N=3 for pathogen-exposed cells) combined with requiring peptides to appear in at least two out of three replicates effectively filters artifacts. Furthermore, downstream validation with binding prediction algorithms (netMHCpan/netMHCIIpan) and experimental confirmation in exposed cells provides multiple layers of verification, ensuring that identified peptides are genuine MHC-presented antigens rather than contaminants.

The accessibility of our approach is a key advantage: researchers can use our published common binding motif data and bioinformatic tools to query pathogen proteomes in an afternoon, with no requirement for expensive wet-lab validation unless desired. When wet-lab validation is performed, our results demonstrate that even three cell lines can be sufficient to experimentally confirm antigenic regions, as we showed with *S. enterica*.

### Practical Implementation and Broad Applicability

Our workflow is designed to be both rigorous and practical. We selected 30 cell lines to create a panel representing the world’s most common MHC genetic variants, achieving coverage of approximately 75% of the most common global HLA alleles. Based on the common sequence motifs that 24 of these 30 cell lines can present, we designed a bioinformatic tool that predicts globally presentable immunogenic regions of interest (ROIs) from a pathogen’s proteome. This method can be applied to any pathogen—whether novel emerging threats requiring rapid vaccine development, or established pathogens where previous vaccine attempts have failed due to various reasons that may include inadequate consideration of HLA diversity.

Our workflow identifies 9-mer peptides that are presentable by diverse MHCs; however, selecting broader regions of the target protein in downstream analysis ensures the inclusion of all globally presentable 9-mers within those regions. As observed in our *Salmonella enterica* (Figure 5 and 6) and SARS-CoV-2 Spike protein (Figure 7) analyses, ROIs are typically characterized by clusters of overlapping presentable peptides rather than isolated 9-mers. The next validation step confirms that predicted regions are presentable by MHC molecules through immunopeptidomics on one or a few cell lines—as we demonstrated, substantial validation can be achieved with minimal wet-bench work. The final step tests whether presented antigens are truly immunogenic by determining if they are recognized by MHC- allele-matched T-lymphocytes using ELISPOT assays.

**Figure 5.**
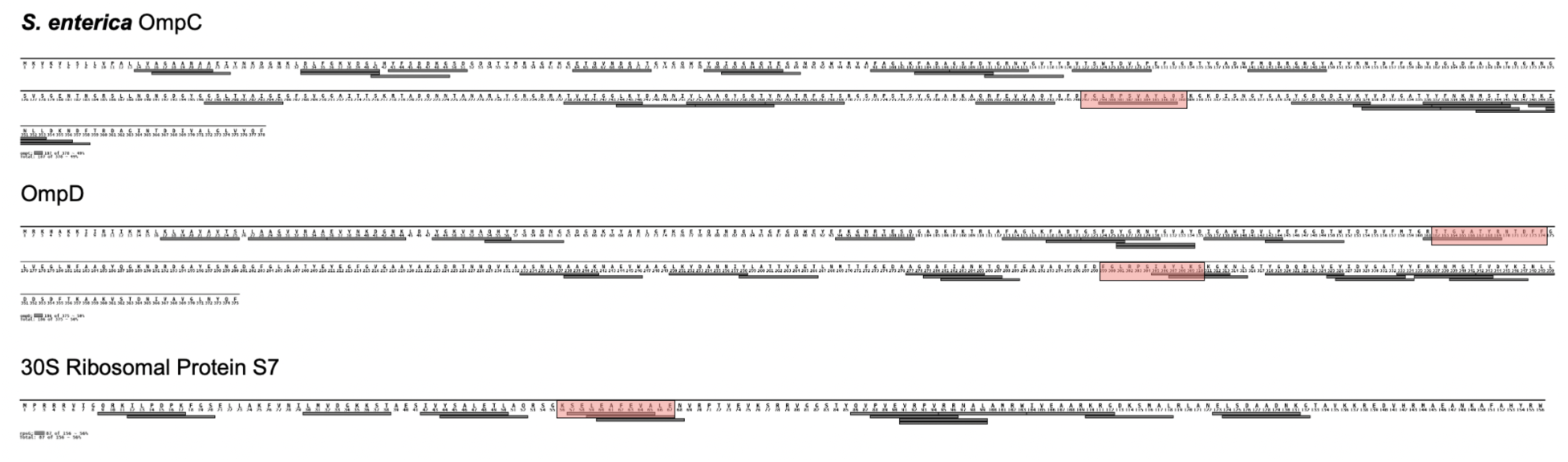
*S. enterica* 9-mers predicted to be globally presentable based on sequence similarity to one of the eighteen common binding motifs cross matched with highly presented immunopeptidomic data acquired by MAE-LC-MS/MS to produce regions of the protein that are likely to be globally immunogenic. Overlapping sequences shown of the OmpC, OmpD, and 30S ribosomal proteins detected as MHC-presented after a 24-h bacteria lysate exposure. Each epitope was considered a positive ID if it was found in a minimum of two out of three replicates. Grey bars = predicted ROIs. Highlighted bars = detected epitopes.

**Figure 6.**
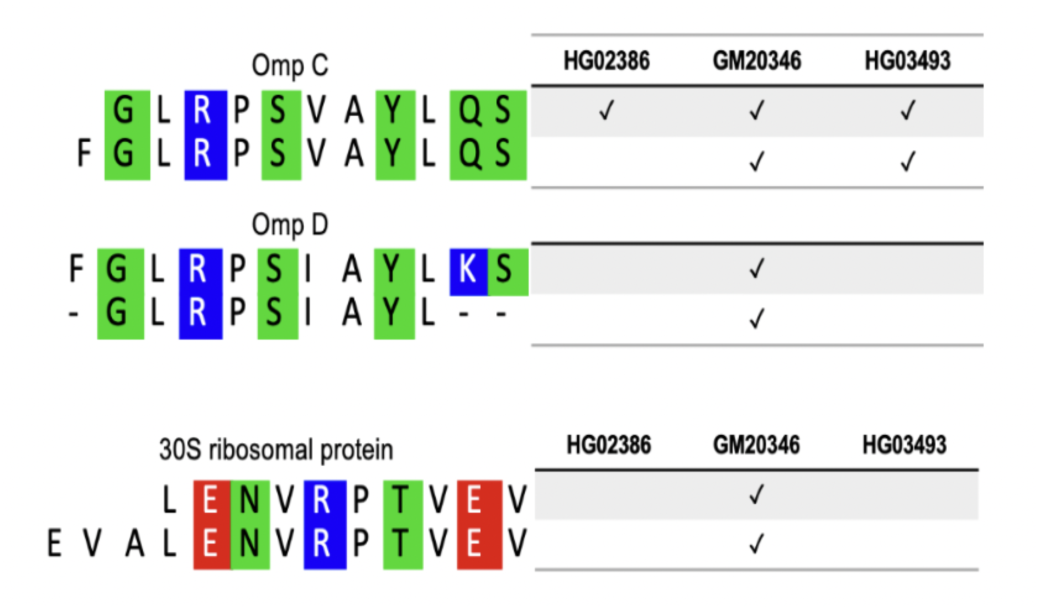
Overlapping sequences shown of the OmpC, OmpD, and 30S ribosomal proteins detected as MHC-presented after a 24-h bacteria lysate exposure. Each epitope was considered a positive ID if it was found in a minimum of two out of three replicates.

**Figure 7.**
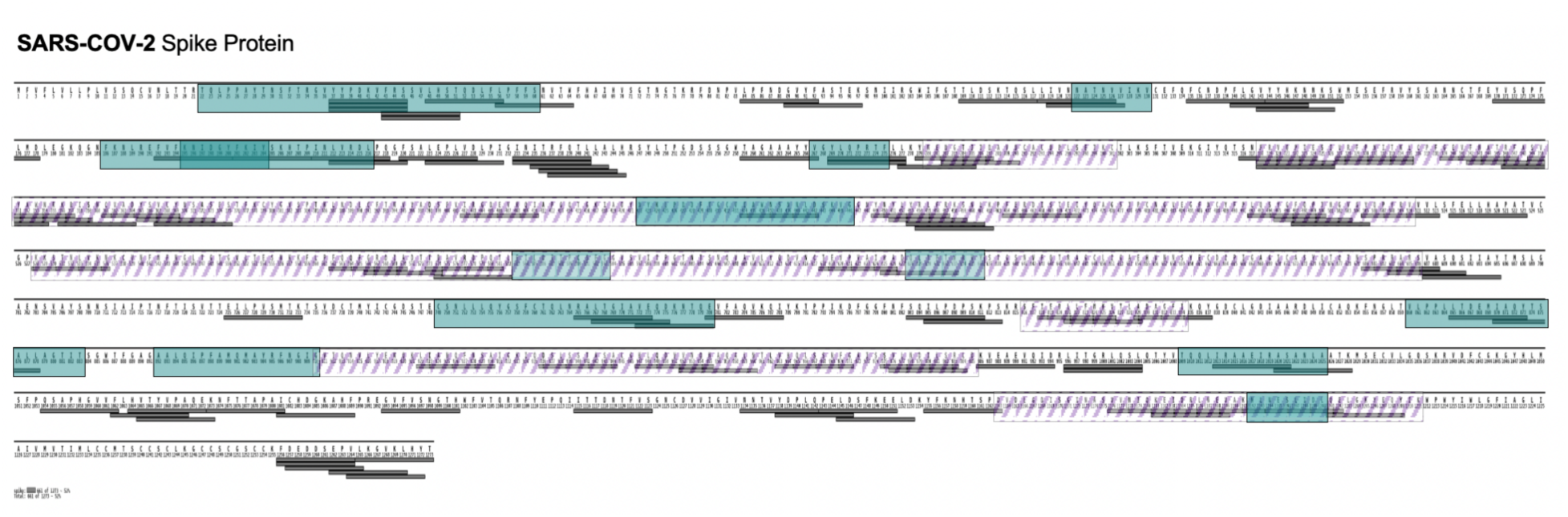
SARS-CoV-2 Spike protein 9-mers predicted to be globally presentable based on sequence similarity to one of the eighteen common binding motifs cross matched with literature identified antigenic regions. Blue regions are highly presented HLA-I and -II presented immunopeptides identified by IP-LC-MS/MS (Weingarten-Gabbay *et al*., 2021, 2024). The diagonal striped regions are regions of the protein that have been identified as antigenic based on the structure of the fragment.

It is important to note that although many of the detected *S. enterica* peptides aligned with our predicted immunogenic regions, not every presented antigen will have a cognate T-cell receptor or will induce a T- cell response. A peptide can be tested if it is immunogenic by determining if it is recognized by an MHC allele matched T-lymphocyte. An Enzyme-Linked ImmunoSpot (ELISPOT) sandwich enzyme-linked immunosorbent (ELISA) assay can be used to test if each peptide is immunogenic. Our experimental conditions—exposing cells to S. enterica lysate rather than purified proteins—likely explain why MAE yielded only one or two peptides per protein while our bioinformatic mapping predicted more extensive coverage. If cells had been exposed to individual purified proteins, we expect that more MAE-detected peptides would align with bioinformatically predicted regions. This limitation does not diminish the value of our predictions; rather, it highlights that the bioinformatic predictions cast a wide net that requires experimental validation to prioritize the most promising candidates. Given unlimited resources, all predicted ROIs could be tested both in diverse cell line panels and in animal models to comprehensively characterize their immunogenicity.

Additionally, while our 30-cell line panel represents 75% of the world’s most common HLA alleles, rare alleles and population-specific variants may require supplemental testing. However, by capturing the majority of global HLA diversity, our approach ensures broad population coverage while remaining practically feasible for most vaccine development programs.

### Validation with Established Antigens

Our validation approach leveraged pathogens with well-characterized antigenic regions to demonstrate that our method successfully identifies known immunogenic targets. For *S. enterica* serovar Typhimurium, we detected OmpC in all three cell lines tested, along with nested peptide sets from OmpD and 30S ribosomal protein S7—patterns indicative of highly immunogenic antigens. Importantly, exposure to Omp proteins has been shown to protect hosts against lethal doses of S. enterica serovar Typhimurium with 100% efficacy, and both OmpC and OmpD are known to be highly immunogenic porins in S. enterica serovar Typhi. All experimentally identified epitopes showed substantial overlap with our bioinformatically derived regions of interest, supporting their potential as broadly immunogenic vaccine candidates. Similarly, for the SARS-CoV-2 Spike protein, most presented peptides identified in recent immunopeptidomics studies occurred in regions our method predicted to contain multiple presentable 9- mers across common binding motifs. Additional characterized antigenic regions—including a putative superantigen, the receptor-binding domain, the fusion peptide, and heptad repeats—also overlapped with our globally presentable predictions. These validations demonstrate that our method successfully captures established antigenic regions while simultaneously predicting which regions are likely to be broadly immunogenic across diverse populations.

### Future Directions and Clinical Translation

Mouse model testing with our S. enterica candidates is currently underway, representing the next step toward clinical validation. The method can readily be applied to neglected tropical diseases and pandemic preparedness efforts, where rapid identification of broadly protective antigens is critical. While our panel was derived from the 1000 Genomes Project—the most comprehensive publicly available resource with both genetic information and immortalized cell lines representing global diversity—it could be supplemented with additional cell lines targeting specific populations or rare HLA alleles if researchers identify particular genetic backgrounds requiring additional coverage.

The most significant barrier to widespread adoption of our approach is not technical or regulatory, but rather awareness. This methodology could become standard practice in early vaccine development, serving as a screening tool to ensure that antigens with broad population coverage advance to clinical trials. Such an approach would naturally encourage more diverse participant enrollment in Phase I/II trials, as researchers would have greater confidence that their vaccine candidates should work across multiple genetic backgrounds. This represents a fundamental shift from current practice, where diversity is considered primarily as a trial enrollment issue rather than as a core principle guiding antigen selection from the earliest stages.

## Conclusion

We have developed a globally representative immunopeptidomics approach that addresses a critical equity gap in vaccine development. By considering HLA diversity at the earliest stages of antigen discovery rather than after clinical trial failures, this method has the potential to increase vaccine success rates while ensuring that new vaccines provide broad protection across diverse populations. The combination of computational prediction with experimental validation using a diverse cell line panel offers a practical, accessible pathway for researchers developing vaccines against any pathogen. As the field moves toward more equitable and inclusive vaccine development, approaches like ours that embed diversity considerations into the fundamental research process will be essential for creating vaccines that truly protect global populations.

