## Supplementary Data for "A Globally Representative Immunopeptidomics Approach to Identify Population-Wide Vaccine Candidates"

**Supplemental Data:**

**Table S1 – MHC Class I (HLA A, HLA-B, and HLA-C) haplotype data for the 30 selected 1000 Genomes Project cell lines** (Auton, 2015)**.** Data represent global genetic diversity for an in-house pre-clinical trial cell line panel assigned with 30× whole-genome NGS by xHLA (Xie et al. 2017).
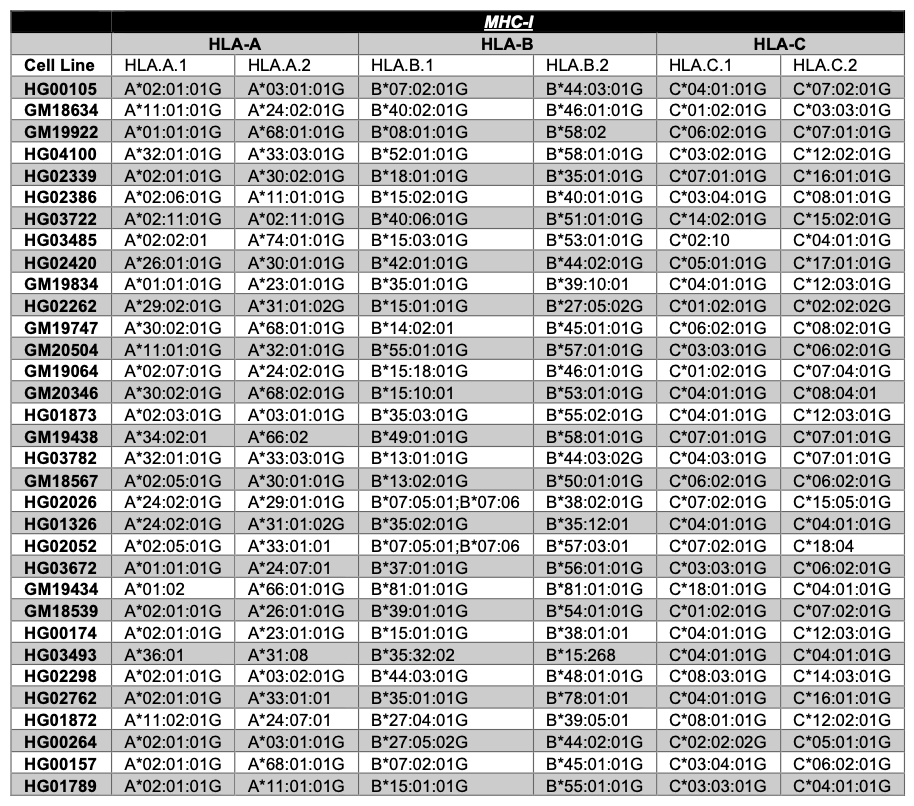

**Table S2 – MHC Class II (HLA-DPA,DPB, DQA, and DQB) haplotype data for the 30 selected 1000 Genomes Project cell lines** (Auton, 2015)**.** Data represent global genetic diversity for an in-house pre-clinical trial cell line panel assigned with 30× whole-genome NGS by xHLA (Xie et al. 2017).

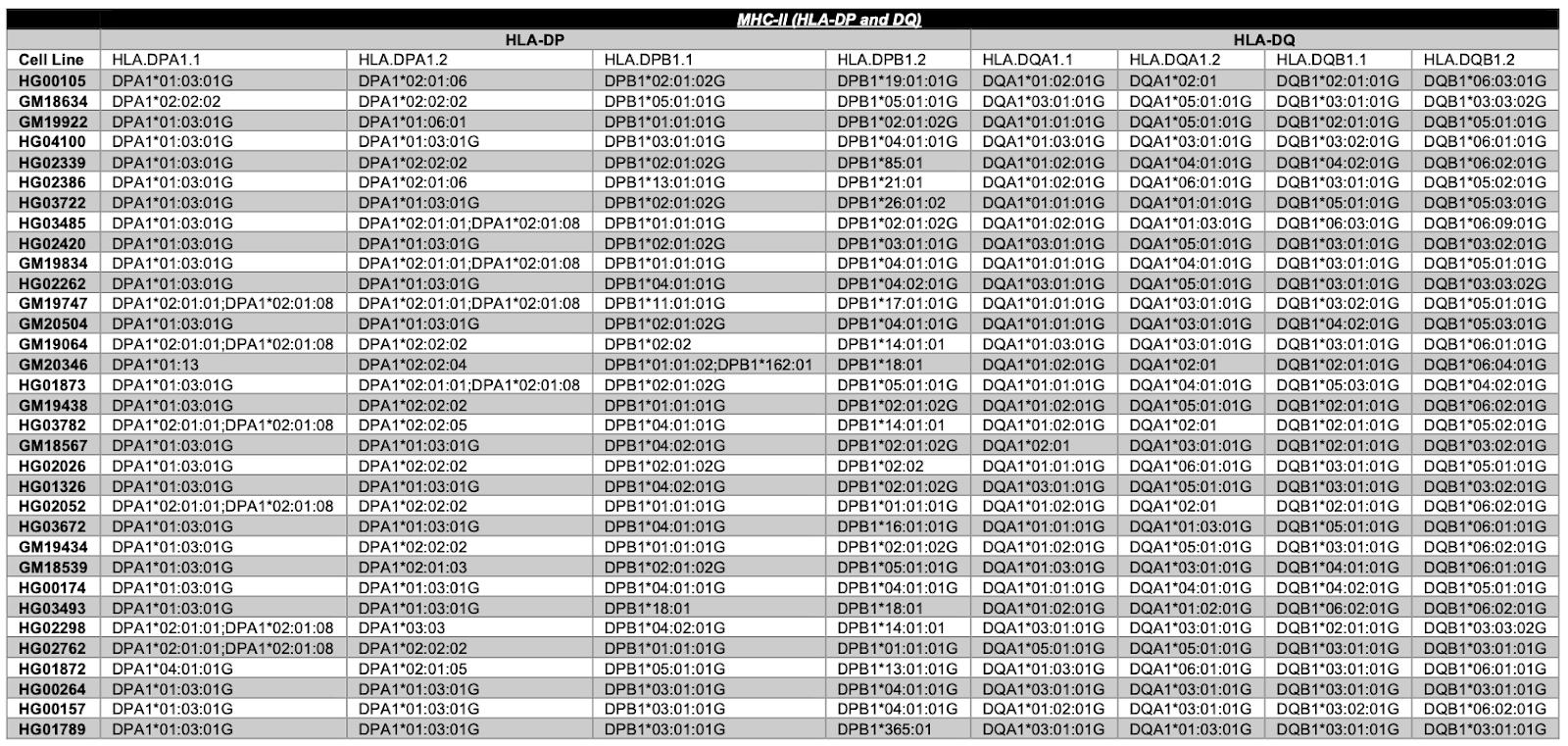

**Table S3 – MHC Class II (HLA-DRB1, DRB3, and DRB4) haplotype data for the 30 selected 1000 Genomes Project cell lines** (Auton, 2015)**.** Data represent global genetic diversity for an in-house pre-clinical trial cell line panel assigned with 30× whole-genome NGS data by xHLA (Xie et al. 2017).

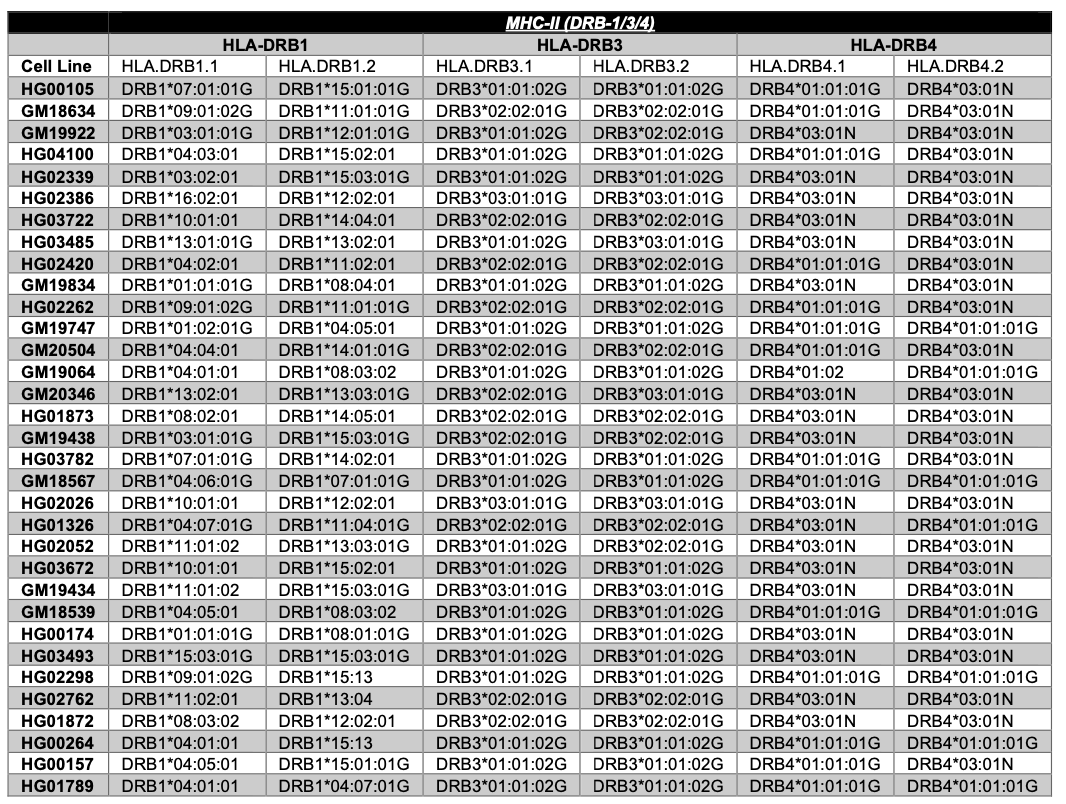

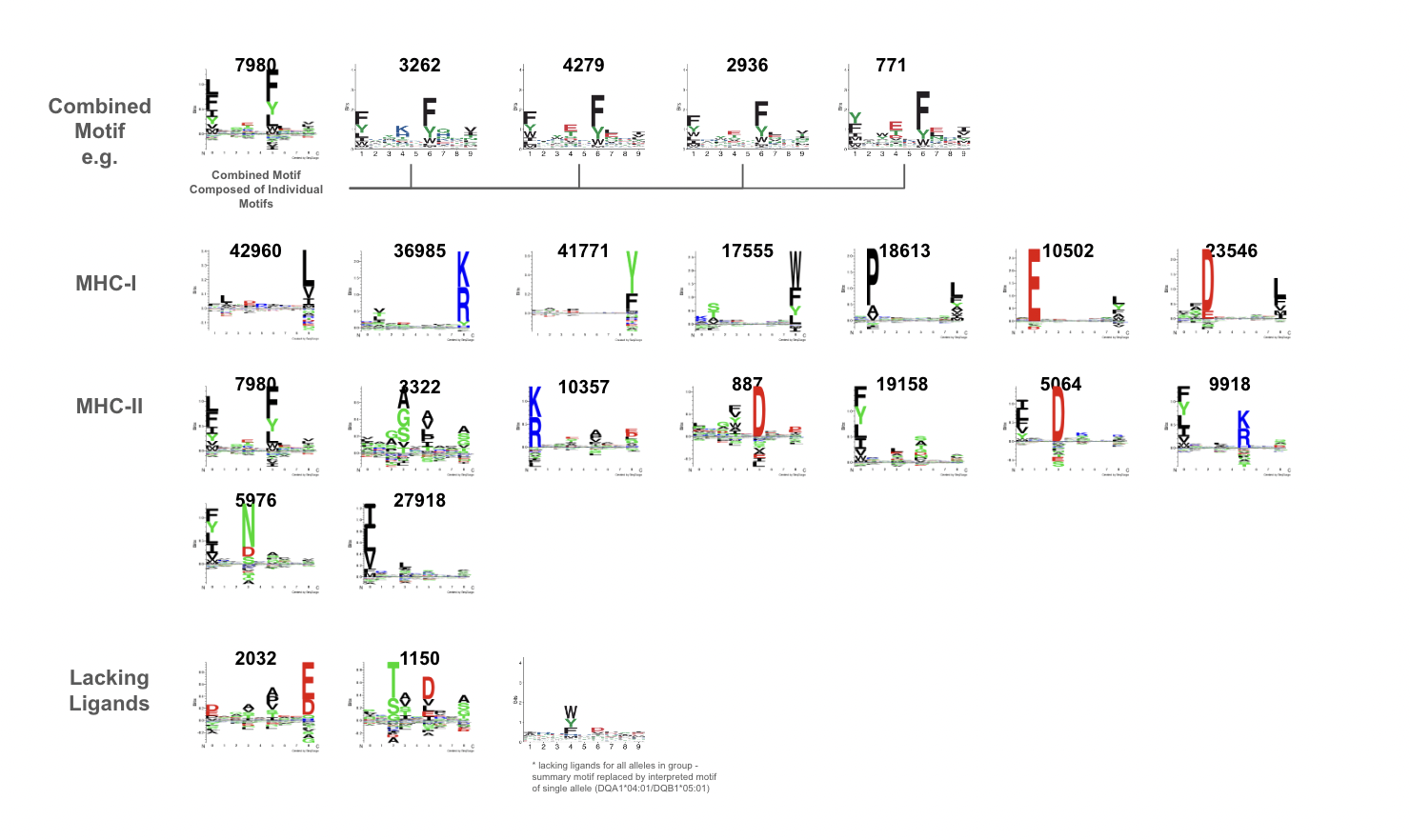

Figure S1. Sequence logo plots of common binding motifs based on MHC Motif Atlas library peptides from the top fifteen of the 1000 Genomes Project cell line panel (Tadros *et al.*, 2023; Auton, 2015). Common motifs needed to occur in at least 5 cell lines to be included. Due to a lack of downloadable ligands for 3 common motifs, the groups were omitted as available peptides were not representative of the group. A total of 7 MHC-I and 9 MHC-II common motifs were identified.

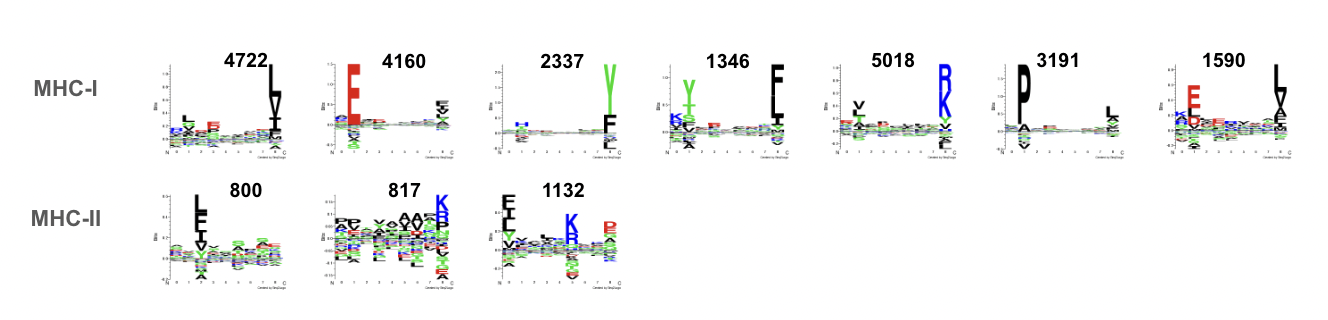

Figure S2. Sequence logo plots of common binding motifs found in the fourteen of the 1000 Genomes Project cell lines (Auton, 2015). Common motifs from mild acid elution data of fourteen randomly selected cell lines. Common motifs needed to occur more than once to be grouped and included. A total of 7 MHC-I and 3 MHC-II common motifs were identified.

**Table S4 – *Salmonella enterica* immunopeptides identified and quantified eluted from both MHC-I and -II by MAE.** LC-MS/MS LFQ intensity values of bacteria-derived MHC-presented epitopes, grouped by protein, from three 1000 Genome Project (Auton, 2015) B-lymphocytes exposed with a bacteria lysate for 24-h and their paired plain MAE controls. Epitope quantification was performed by taking the average value of the individual LFQ values of all the peptides in a given epitope. Since many epitopes are singlets (epitopes with a single peptide, as noted in the peptide per epitope (P/E) column) the average intensity value of all the replicates for the singlet epitope is reported. The *S. enterica* database included both Swiss-Prot (reviewed/annotated experimental data) and TrEMBL (unreviewed/interpreted from genomic information) proteins (SP/Tr column).

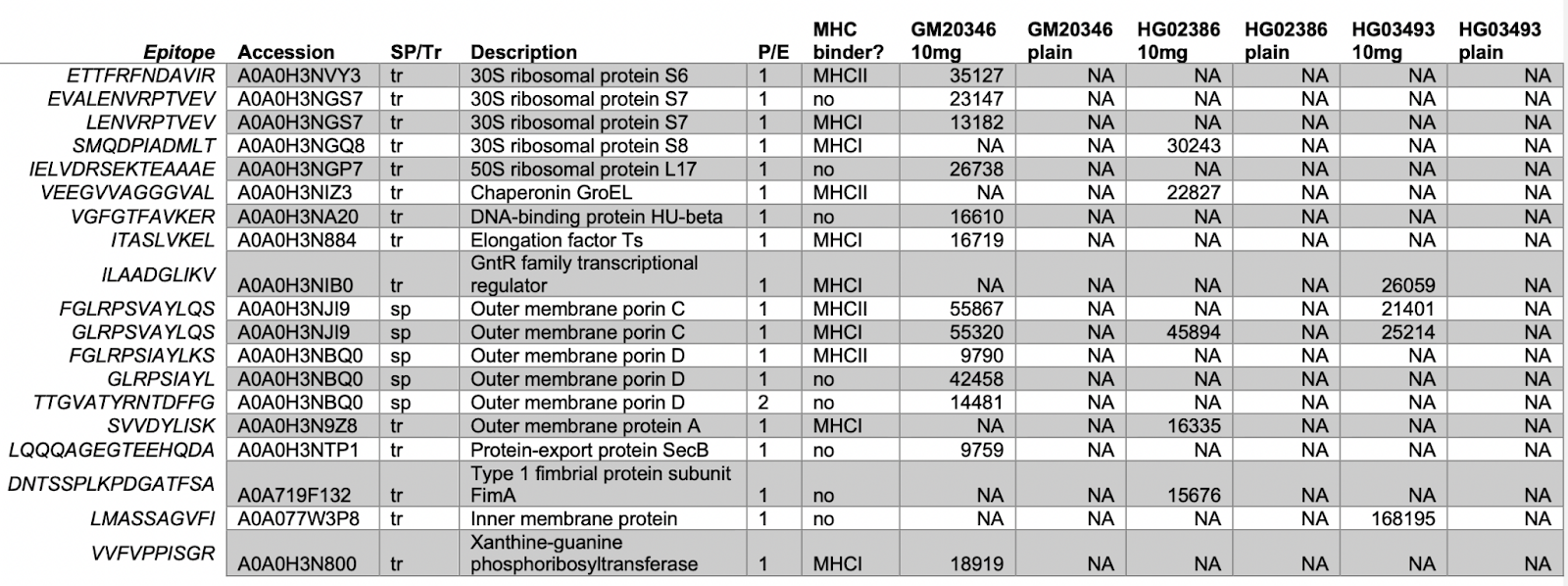

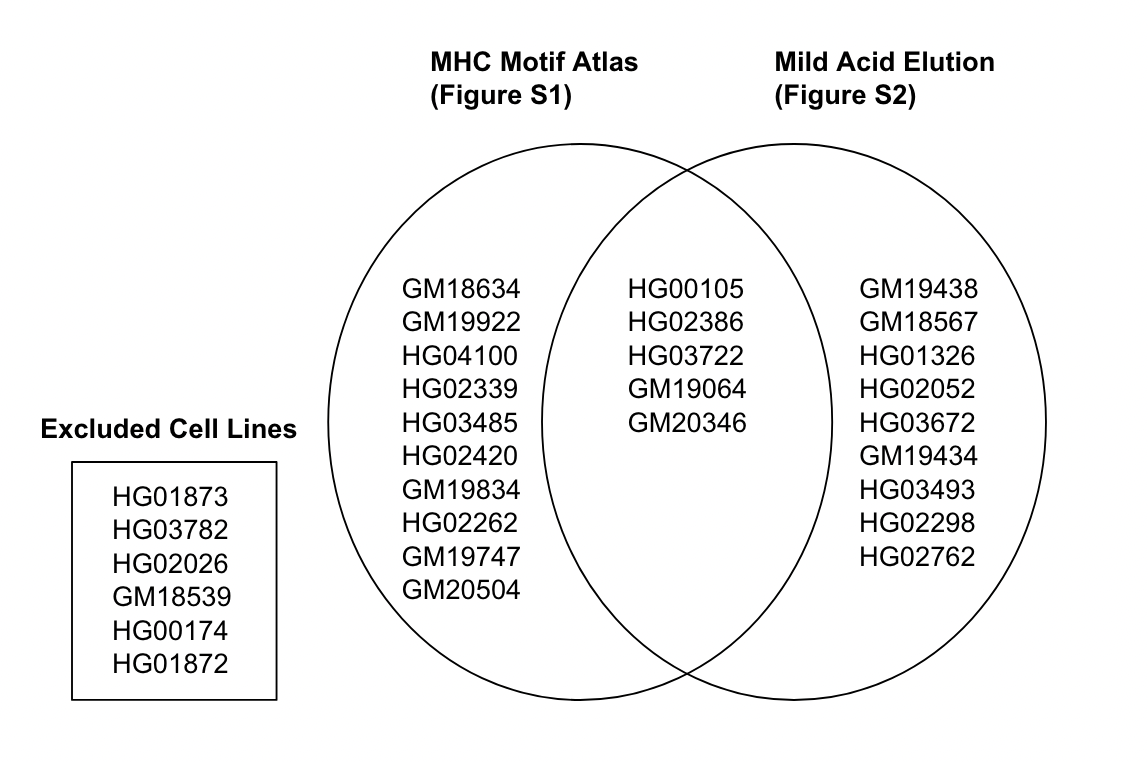

Figure S3. List of Specific Cell Lines Used for Generating Common Binding Motifs

**Table S5 – SARS-COV-2 Spike Protein Region of Interest**

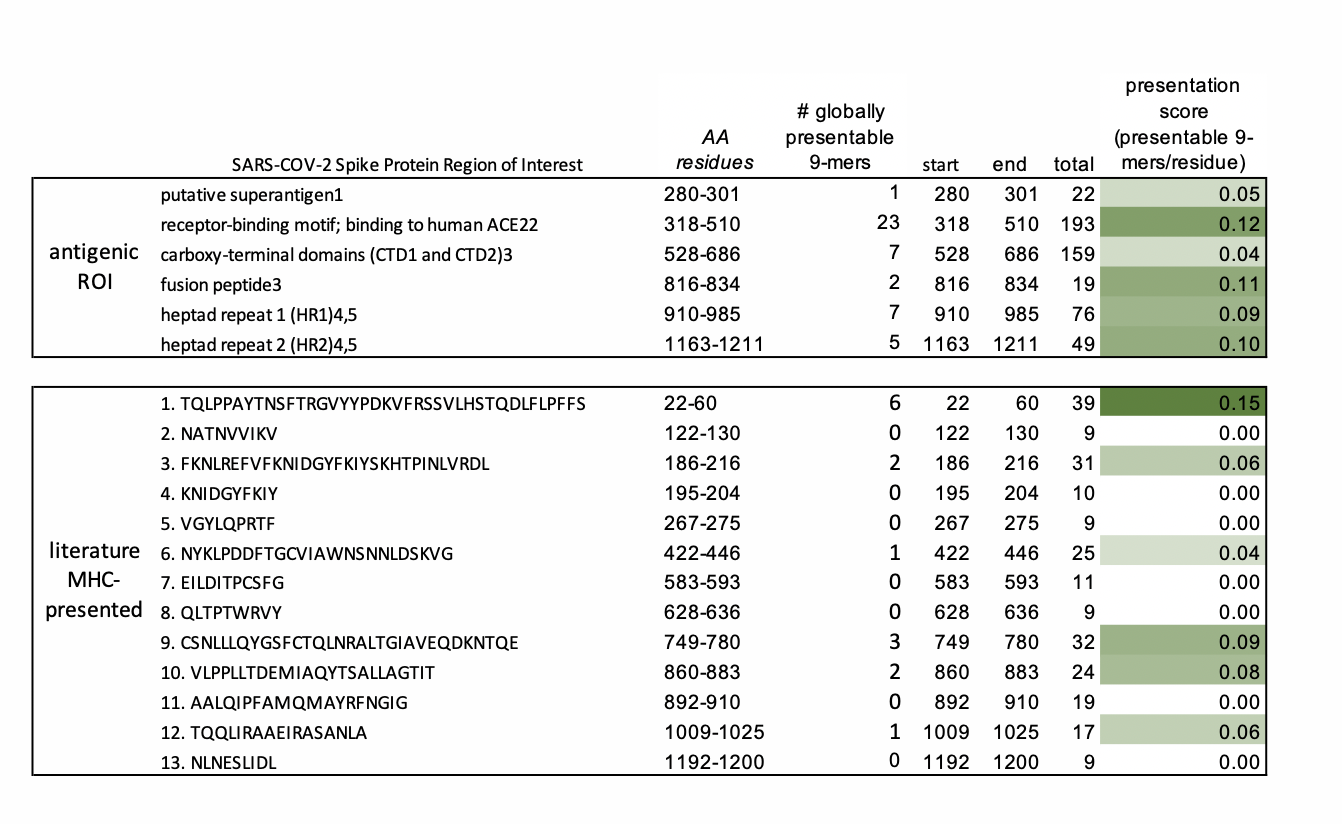
